# Systematic analysis of the type VII secretion system in *Streptococcus gallolyticus* subsp. *gallolyticus* reveals genomic diversity and functional associations

**DOI:** 10.64898/2026.04.05.716583

**Authors:** Guadalupe Calderon, Jyoti Tamang, Sarah Woodfin, Isaac Prah, Julian Hurdle, Yi Xu

## Abstract

*Streptococcus gallolyticus* subsp. *gallolyticus* (*Sgg*) is an opportunistic pathobiont associated with bacteremia, infective endocarditis, and colorectal cancer. However, the genomic diversity of this subspecies and the distribution of key virulence determinants, particularly the type VII secretion system (T7SS), remain poorly characterized. Here, we performed genomic analyses of 76 *Sgg* strains from diverse geographic and host origins. Core- and pan-genome analyses, multi locus sequence typing, and phylogenetic reconstruction revealed dominant sequence types (STs) that correlate with geographic origin or source of isolation. Furthermore, systematic characterization of the T7SS locus identified five new T7SS subtypes and demonstrated a strong association between T7SS subtype and ST. We further expanded the known repertoire of T7SS LXG domain-containing polymorphic toxins (LXG toxins) in *Sgg* substantially through genome-wide searches. Distinct distribution patterns were observed for the LXG toxins across the strains. Lastly, our data indicated that T7SS subtype was significantly associated with biofilm formation capacity of *Sgg* strains. Together, these findings advance our understanding of *Sgg* genomic diversity, reveal substantial lineage-associated variation in T7SS architecture and effector repertoires, and suggest a previously unrecognized connection between T7SS and biofilm formation in *Sgg*.

## 1 Introduction

*Streptococcus gallolyticus* subspecies *gallolyticus* (*Sgg*), previously known as *Streptococcus bovis* biotype I, is a commensal bacterium of the gastrointestinal tract of various animals, including ruminants, marsupials, and birds. In humans, it is estimated to be present in approximately 5 – 30% of healthy individuals (Jans and Boleij, 2018; Pasquereau-Kotula *et al*., 2018; Périchon *et al*., 2022). In addition to its commensal lifestyle, *Sgg* acts as an opportunistic pathogen that causes bacteremia and infective endocarditis (IE), accounting for approximately 11-14% of IE cases worldwide (Hoen *et al*., 2005; Corredoira *et al*., 2008; Vogkou *et al*., 2016). A distinctive feature of *Sgg*-associated IE and bacteremia is their strong association with colorectal cancer (CRC). This relationship has been extensively documented in numerous case reports and cohort studies over several decades, with an average reported incidence of approximately 60% (Gupta, Madani and Mukhtar, 2010; Abdulamir, Hafidh and Abu Bakar, 2011; Boleij *et al*., 2011a; Corredoira *et al*., 2015; Pasquereau-Kotula *et al*., 2018). Several studies have also reported preferential enrichment of *Sgg* in tumor-bearing colons compared with normal colons (Abdulamir, Hafidh and Bakar, 2010; Kumar *et al*., 2017; Aymeric *et al*., 2018; Thomas *et al*., 2019; Périchon *et al*., 2022; Wang *et al*., 2025). Furthermore, in multiple animal models of CRC, oral gavage of *Sgg* resulted in larger tumors, increased tumor burden, and activation of cancer-related pathways (Kumar *et al*., 2017, 2018; Zhang *et al*., 2018; Pasquereau-Kotula *et al*., 2023a; Wang *et al*., 2025), suggesting that *Sgg* actively promotes colon tumor development. At the cellular level, *Sgg* has been shown to adhere to colon epithelial cells, stimulate colon cancer cell proliferation, activate cancer-related and stress response pathways, and translocate across a monolayer of epithelial cells (Boleij *et al*., 2011b; Kumar *et al*., 2018; Martins *et al*., 2020; Taddese *et al*., 2020; Pasquereau-Kotula *et al*., 2023b). Despite these advances, the molecular mechanisms and bacterial factors that mediate *Sgg* virulence and host interactions remain incompletely understood.

Genomic and phenotypic heterogeneity is common among many species of pathogenic bacteria. Efforts to define strain lineages, sequence types (STs), core and accessory genomes, and correlations between genotypes and phenotypes have provided important insights for tracking and preventing the spread of infectious diseases. They have also facilitated the identification of common or lineage-specific biomarkers and virulence factors that inform molecular epidemiological surveillance, diagnosis, treatment strategies, and mechanistic studies (Schürch and Siezen, 2010; Ladner *et al*., 2019). With regard to *Sgg*, multiple studies have demonstrated considerable variation in virulence phenotypes and host interactions among strains (Sillanpää *et al*., 2008; Vollmer *et al*., 2010; Grimm *et al*., 2017a, 2017b, 2018; Kumar *et al*., 2018; Taddese *et al*., 2020; Pasquereau-Kotula *et al*., 2023a). Several studies have attempted to type *Sgg* strains and characterize their genomes, primarily using isolates of European origin (Dumke *et al*., 2014; Shibata *et al*., 2014; Périchon *et al*., 2025). These studies highlighted a complex genomic landscape in *Sgg*; however, further investigation is needed to better understand *Sgg* lineages and the genetic factors contributing to the phenotypic differences.

The type VII secretion system (T7SS) is a specialized secretion system present in Gram-positive actinobacteria (T7SSa) and firmicutes (T7SSb). T7SS-secreted effector proteins have been shown to play roles in interbacterial competition, interactions with the host, and nutrient acquisition. Accordingly, they are often crucial for survival under niche conditions, host colonization, and virulence (Unnikrishnan *et al*., 2017; Rivera-Calzada *et al*., 2021; Spencer and Doran, 2022; Garrett and Palmer, 2024). Two recent studies shed light on the importance of T7SS to *Sgg* virulence and host colonization. A T7SS from *Sgg* strain TX20005 was demonstrated to be critical for virulence (Taylor *et al*., 2021). Deletion of the T7SS abolished the ability of the strain to stimulate CRC cell proliferation and reduced bacterial adherence to colonic epithelial cells. *In vivo*, T7SS deletion resulted in significantly reduced bacterial colonization of the colon and impaired the ability of the strain to promote colon tumor development. In addition, Teh and colleagues characterized TelE, a T7SS-dependent LXG domain-containing polymorphic toxin (LXG toxin) from *Sgg* strain UCN34, and demonstrated that this toxin exhibits pore-forming toxicity against *E. coli* (Teh *et al*., 2023). Together, these findings support the T7SS as an important virulence determinant for *Sgg* that warrants further investigation.

The T7SS is known to display substantial heterogeneity between bacterial species and among strains within the same species (Kneuper *et al*., 2014; Warne *et al*., 2016; Jäger, Kneuper and Palmer, 2018; Lebeurre *et al*., 2019; Zhou *et al*., 2022; Spencer *et al*., 2023). This heterogeneity is thought to contribute to specialized virulence traits and niche adaptation (Kneuper *et al*., 2014; Jäger, Kneuper and Palmer, 2018; Spencer *et al*., 2021). Previous work by Périchon and colleagues has described two highly divergent T7SS subtypes within *Sgg*; subtype 1, represented by the T7SS in strain UCN34 and subtype 2, represented by TX20005 (Périchon *et al*., 2025). These two subtypes exhibit extensive differences in sequence identity as well as in the genetic organization of genes encoding core components of the T7SS. The C-terminal region of EssC, an ATPase essential for the secretion activity, has been demonstrated to be a specificity determinant of T7SSb secretion (Warne *et al*., 2016; Jäger, Kneuper and Palmer, 2018; Lebeurre *et al*., 2019; Zhou *et al*., 2022; Spencer *et al*., 2023). However, a systematic investigation of the T7SS in *Sgg* particularly with respect to variation in the EssC C-terminal region, effectors, and their associations with broader genomic features and functional traits, has not been conducted.

In this study, we carried out a comprehensive genomic characterization of 76 *Sgg* strains isolated from human and animal sources from diverse geographical locations. Multi locus sequence typing (MLST), core- and pan-genome analyses, and phylogenetic tree construction were performed. Furthermore, we systematically examined the T7SS locus and the repertoire of LXG toxins across the 76 genomes. These analyses uncovered dominant sequence types (ST), new T7SS subtypes, and novel LXG toxins. We further performed functional assays with a cohort of *Sgg* strains to evaluate two virulence traits, biofilm formation and stimulation of CRC cell proliferation. More importantly, correlation analysis revealed significant associations among ST, isolation source, T7SS subtype, and biofilm formation capacity. Together, these results provide new insights into the relationship between genomic diversity, T7SS heterogeneity, and virulence phenotypes in *Sgg*, thereby advancing our understanding of its genomic landscape and virulence-associated features.

## 2 Materials and Methods

### 2.1 Cells, bacterial strains, and culture conditions

Human colon cancer cell lines HT29 and HCT116 were obtained from ATCC and cultured in Dulbecco’s Modified Eagle Medium (DMEM) with Hams F-12 50:50 Mix [+] L-glutamine, 15 mM HEPES (Corning) supplemented with 10% fetal bovine serum (FBS) at 37°C in a humidified chamber with 5% CO2. *Sgg* strains in the lab collection were primarily a gift from Dr. Barbara E. Murray, University of Texas Health Science Center, Houston, Texas. They were originally collected from the blood of patients with IE and/or bacteremia from two different hospitals in the United States (Sillanpää *et al*., 2008) (Supplementary Data, Table S2). Strain UCN34 was provided by Dr. Shaynoor Dramsi, Institut Pasteur, Paris, France. Strains BAA-2069 and ATCC 43143 were purchased from ATCC. *Sgg* strains were cultured in brain heart infusion broth at 37°C with shaking or grown on tryptic soy or BHI agar plates at 37°C overnight.

### 2.2 Genome sequencing, assembly, and annotation

Genomic DNA was extracted from *Sgg* using Qiagen’s QIAamp DNA Mini Kit. Whole genome sequencing was performed at SeqCenter (Pittsburg, PA) via a combination of Illumina short-read sequencing and Oxford Nanopore long-read sequencing. Raw reads were filtered through FastP or FastPLong (Chen *et al*., 2018) for read quality and trimming processing. The cleaned-up short and long reads were used for hybrid assembly using Unicycler (Wick *et al*., 2017). Genomes were annotated using Prokka (Seemann, 2014).

### 2.3 Acquisition of *Sgg* genomes from the NCBI

The National Center for Biotechnology Information (NCBI) Genomes database was searched using the search term “*Streptococcus gallolyticus”* and the resulting genomes (available up to July 31, 2025) were downloaded. As *S. gallolyticus* consists of several subspecies, we used the online tool Type (Strain) Genome Server (TYGS) (https://tygs.dsmz.de/) (Meier-Kolthoff and Göker, 2019) to filter out genomes of other subspecies. The confirmed *Sgg* genomes were then assessed for assembly quality. Only genomes with an N50 larger than 100 kb and contain less than 50 contigs were kept for subsequent analyses. This resulted in 46 *Sgg* genomes from NCBI. For consistency, these genomes were also annotated using Prokka.

### 2.4 General characterizations of the genomes

MLST for *Sgg* genomes was carried out using MLST 2.0 at the Center for Genomic Epidemiology (Camacho *et al*., 2009; Larsen *et al*., 2012) (https://cge.food.dtu.dk/services/MLST/) and PubMLST (https://pubmlst.org/) (Jolley, Bray and Maiden, 2018), following the typing scheme developed previously for *Sgg* using seven housekeeping genes (*aroE*, *glgB*, *nifS*, *p20*, *tkt*, *trpD*, and *uvrA*) (Dumke *et al*., 2014). To search for virulence factors in the genomes, the Virulence Factor Database (VFDB) (https://www.mgc.ac.cn/cgi-bin/VFs/v5/main.cgi) (Chen *et al*., 2005) was used. The Resistance Gene Identifier (RGI) (https://card.mcmaster.ca/analyze/rgi) (Alcock *et al*., 2023) was used to search for potential genes involved in resistance to antimicrobial agents.

### 2.5 Core- and pan-genome analysis and phylogenetic tree construction

Roary (Page *et al*., 2015) was used to analyze core- and pan-genomes. Core genes are defined as present in ≥ 99% strains. Proteins with a minimum of 90% amino acid sequence identity over most of the length are considered the same. PRANK was used to create a multiple sequence alignment of core genes (Löytynoja and Goldman, 2010). RAxML-NG (Kozlov *et al*., 2019) was used to build a phylogenetic tree based on maximum likelihood and bootstrap values were calculated based on 100 bootstrap replicates. The best model was then used to visualize and annotate the phylogenetic tree using iTOL (Letunic and Bork, 2007).

### 2.6 T7SS identification, classification of subtypes, and identification of putative LXG toxins

To identify the T7SS locus within each genome, a custom BLAST database of all 76 *Sgg* genomes was created using the makeblastdb command. This database was then searched for sequence homology to EsxA and EssC, the two most conserved T7SS core components, using tblastx. Strains showing significant homology to these proteins were then examined for the presence of genes encoding other core components of the T7SS secretion machinery (*esaA*, *essA*, *esaB* and *essB*) as well as genes downstream of the core machinery genes. The T7SS was further classified into subtypes based on the variable C-terminal 200 amino acid sequence of EssC and gene order downstream of *essC*.

To identify putative LXG toxins, the database was searched using Blastp with queries including characterized LXG toxins and an LXG domain consensus sequence (pfam04740) from NCBI-Conserved Domains Database (CDD). A minimum of 90% amino acid sequence identity over most of the length is used as a cut off for being considered the same LXG protein. Motif Search at GenomeNet (https://www.genome.jp/tools/motif/) was used to identify protein motifs in the putative LXG toxins against Pfam, NCBI-CDD, and Prosite (E value cut off 0.001).

### 2.7 Bacterial biofilm assay

Biofilm assay was carried out as described previously (Grossman et al., 2021) with slight modifications. Briefly, overnight cultures of *Sgg* were diluted to an OD600 of 0.1 in BHI broth. The bacterial suspension or BHI (media control) were added to the wells in a 96-well plate and incubated at 37°C for 24 hours. The wells were then washed, stained with crystal violet, washed again and incubated with methanol. Absorbance at 590nm was then measured.

### 2.8 Cell proliferation assay

This was performed as described previously (Taylor *et al*., 2023) with slight modifications.

Briefly, CRC cells were seeded at a concentration of 1×10^4^ cells per well in a 96-well plate in serum-free DMEM/F-12 (SFM) and incubated overnight. Stationary phase bacteria resuspended in SFM were then added to the cells at an MOI of 1 and incubated with the cells. Bacterial suspensions were also added to wells with no cells to serve as a control for the bacteria only reading. A sublethal dose of trimethoprim (1 μg/mL final concentration) was added at 6 hours of incubation to inhibit bacterial growth. Penicillin (100 units/ml final concentration), streptomycin (100 μg/ml final concentration), and erythromycin (10 μg/mL final concentration) were added at 24 hours and incubated for another 24 hours to eliminate bacteria. The wells were then washed and measured using the cell counting kit (CCK)-8 kit (Apex Bio).

### 2.9 Statistical analysis

One-way ANOVA and Dunnet’s multiple comparisons test was performed for comparison between multiple groups using GraphPad Prism 11. To identify statistical significance of correlations between T7SS subtype, sequence type, isolate source, and geographical origin, *p* values were calculated using permutations chi-square calculation with 10,000 permutations in SciPy (Virtanen *et al*., 2020).

## 3 Results

### 3.1 Genomes of *Sgg* and general features

For our analysis, we used a combination of *Sgg* strains in our lab collection and *Sgg* genomes downloaded from the NCBI Genomes database. The majority of our lab strains have not been sequenced previously. To determine their genome sequence, Illumina shot gun and Oxford nanopore sequencing were performed. Short and long reads were assembled into hybrid assemblies, which resulted in 12 complete genomes and 18 incomplete ones containing 2-15 contigs (Table 1 and Supplementary Data, Table S1). The mean genome coverage for the hybrid assemblies is 574X and ranges from 408-748X. The genome size ranges from 2.2 - 2.5 million bases (Mb), with the mean genome size being 2.3 Mb. The mean GC percentage is 37.5%, and the mean number of coding sequences is 2,220 (Table 1 and Supplementary Data, Table S1).

**Table 1.** Summary of genome features of *Sgg* strains.

| Features | Hybrid Assemblies <sup>a</sup> | NCBI Genomes |
| --- | --- | --- |
| Number of genomes | 30 | 46 |
| Genome coverage (X) | 408 – 748 | 18 - 1193 |
| Mean genome coverage (X) | 574 | 169 |
| Number of contigs | 1 – 15 | 1 – 34 |
| Genome size range (Mb) | 2.2 – 2.5 | 2 - 2.5 |
| Mean genome size (Mb) | 2.3 | 2.3 |
| Mean GC percentage | 37.5 | 37.5 |
| Minimum N50 (kb) | 118 | 113 |
| Mean N50 (Mb) | 0.42 | 1.2 |
| Mean number of coding sequences (CDS) | 2,220 | 2,226 |

<sup>a</sup> Isolates in the lab collection were sequenced using Illumina short-read sequencing and Oxford Nanopore long-read sequencing to create *de-novo* hybrid genome assemblies. Raw reads processing, assembly and annotation was done as described in the Materials and Methods section.

Genomes downloaded from the NCBI Genomes database were further examined to confirm their classification as *Sgg* and to exclude assemblies of poor quality, as described in the Materials and Methods section. This resulted in 46 *Sgg* genomes from NCBI. Among these, 14 are complete genomes, while the remaining contains 2 - 34 contigs (Table 1). The mean genome coverage is 169X and ranges from 18-1193X. The mean genome size, GC percentage, and the number of coding sequences are comparable to those of the genomes generated through our hybrid assemblies (Table 1). Combined with our newly sequenced *Sgg* genomes, this resulted in a total of 76 *Sgg* genomes.

The 76 *Sgg* strains were collected from multiple countries across four continents. The majority of them are clinical isolates with a small number of strains from animal sources (environmental strains). Detailed information on the isolation source, geographic location, and associated clinical conditions of these *Sgg* strains is provided in Supplementary Data, Table S2.

### 3.2 Multi locus sequence typing of *Sgg*

MLST analysis of the geno mes was performed using a previously established *Sgg* typing scheme based on seven housekeeping genes (*aroE*, *glgB*, *nifS*, *p20*, *tkt*, *trpD*, and *uvrA*) (Dumke *et al*., 2014). We were able to determine the ST for 74 strains (Supplementary Data, Table S2 and Fig. 2 below). The remaining two genomes contained assembly gaps in at least one of the seven housekeeping genes, which prevented their assignment to a sequence type. The most prevalent STs are ST12 (11 strains, 14.5%), ST26 (9 strains, 11.8%), ST117 (6 strains, 7.9%), and ST5 (4 strains, 5.3%). The remaining STs are only represented in 1-2 strains. In addition, fourteen new STs (STs 122 – 140) were identified.

The distribution of STs observed here differs from that reported in previous studies (Dumke *et al*., 2014; Périchon *et al*., 2025), which primarily included isolates from France and Germany. This observation suggested a possible geographic-region-related bias. For example, in our analysis, nine of the eleven ST12 strains and all six ST117 strains were isolated in the United States, whereas seven of the nine ST26 strains were from European countries. Correlation analysis indeed revealed a significant correlation between ST and geographic region (*p* = 0.0002) (Table 2).

**Table 2.** Correlations between different traits of *Sgg*.

| Category 1 | Category 2 | p-value<sup>a</sup> |
| --- | --- | --- |
| ST | Geographical region | 0.0002 |
| ST | Isolate source | 0.0002 |
| ST | T7SS subtype | 0.0001 |
| ST | Biofilm | 0.0262 |
| ST | Proliferation | 0.1885 |
| T7SS subtype | Geographical region | 0.0279 |
| T7SS subtype | Isolation source | 0.0119 |
| T7SS subtype | Biofilm | 0.0059 |
| T7SS subtype | Proliferation | 0.4555 |
| Proliferation | Biofilm | 0.5019 |

<sup>a</sup> p-values were calculated using permutations chi-square calculation with 10,000 permutations in SciPy.

We also examined STs among isolates from human vs. environmental sources. Of the four most prevalent STs, ST12, ST117, and ST5 are found in human isolates only. Although the number of environmental isolates was limited, correlation analysis nevertheless showed a significant correlation between ST and isolation source (*p* = 0.0002, Table 2) (Table 2).

### 3.3 Core- and pan-genome analysis, virulence factors, and resistance genes

We determined the core and pangenome of the 76 *Sgg* genomes using Roary (Page *et al*., 2015). Accumulation curves for the number of total and conserved genes vs. the genome number are presented in Fig. 1A and 1B, respectively. The pangenome curve (Fig. 1A) indicates a moderately open pangenome consisting of 7,309 genes, with a Heaps’ law exponent γ of 0.3 (Tettelin *et al*., 2005; Park *et al*., 2019). The core genome curve displays stabilization as the genome number approaches 76 (Fig. 1B). The core genome consists of 1,254 genes, which gives an accessory genome of 6,055 genes. The large accessory genome indicates substantial genomic diversity among *Sgg* strains, consistent with observations in other streptococcal species. The core genes account for approximately 57% of the CDS in an average *Sgg* genome, which falls within the range of what has been reported for other streptococci (Lefébure and Stanhope, 2007; Park *et al*., 2019; Rosconi *et al*., 2022). A core genome alignment was then used to generate a phylogenetic tree of all strains (Fig. 2). Strains with the same ST are clustered together in the tree, as expected.

**Fig 1.**
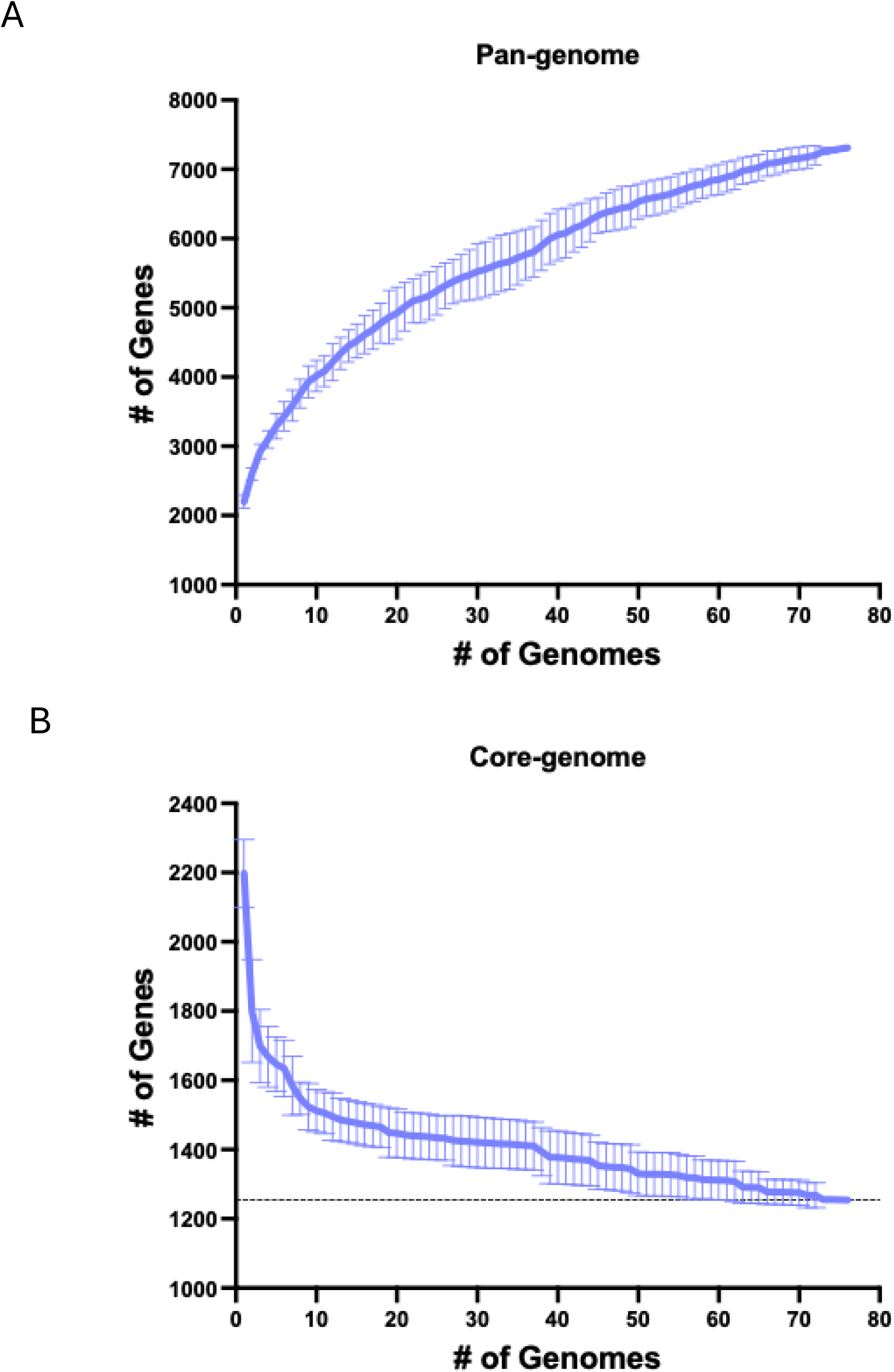
**Pan- and core-genome analysis.** Pan- (**A**) and core-genome (**B**) accumulation curves were determined based on 76 *Sgg* genomes using Roary with 10 permutations. Data shown is the mean ± standard deviation.

**Fig 2.**
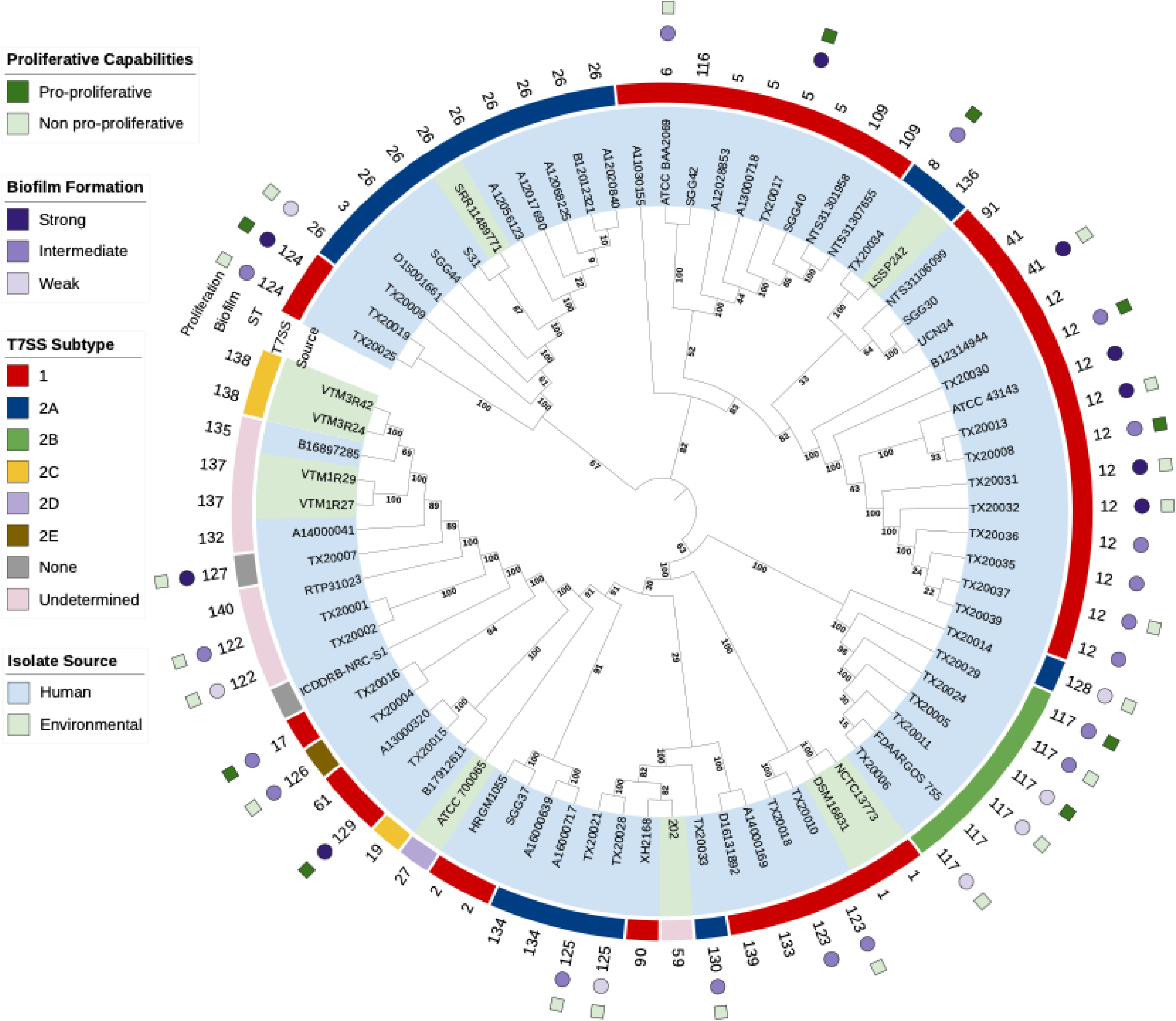
A phylogenetic tree of *Sgg* strains. Multiple sequence alignment of the core-genome genes from 76 strains was performed using PRANK. The alignment was subject to RAxML-NG to generate a phylogenetic tree. The best model was annotated using iTOL. Bootstrap values based on 100 iterations are shown on the branches. Annotations labeled from the outer to the inner part of the circle are: effect on CRC cell proliferation, biofilm formation, sequence type, T7SS subtype, and isolate source, respectively.

We next examined virulence factors encoded by the core and accessory genomes, respectively, by using the online Virulence Factor Database (VFDB) (Chen *et al*., 2005). The majority of core genome-encoded virulence factors are involved in adherence to host proteins (*e.g*., fibronectin binding protein and plasmin receptor), pili assembly (sortase C), and immune evasion or modulation (*e.g*., capsule biosynthesis genes and complement C3-degrading protease) (Table 3). The conservation of these virulence factors in the core genome implies that they are central to the association of this subspecies with its hosts and survival in the host environment. Virulence factors in the accessory genome include additional determinants involved in adherence (*e.g*., pili) and immune evasion or modulation (*e.g*., extra capsule biosynthesis genes and a complement C5a peptidase), further emphasizing host adherence and immune evasion as two potential central features shaping *Sgg*’s lifestyle as a pathobiont. Additionally, factors involved in nutrient acquisition, resistance to bile salts, and fatty acid metabolism, as well as components of T7SS, have also been identified in the accessory genome (Table 3), some of which may contribute to their colonization of the host gastrointestinal tract. The repertoire of virulence factors identified here fall into similar classes as those reported in a recent study by Périchon and colleagues involving primarily French isolates (Périchon *et al*., 2025). We further compared the virulence factors from accessory genomes composed of human and environmental isolates, respectively. We did not identify any virulence factors specific to human isolates. Overall, these findings suggest that the putative virulence factors identified here appear to be distributed across strains from diverse geographic locations or strain sources.

**Table 3.**
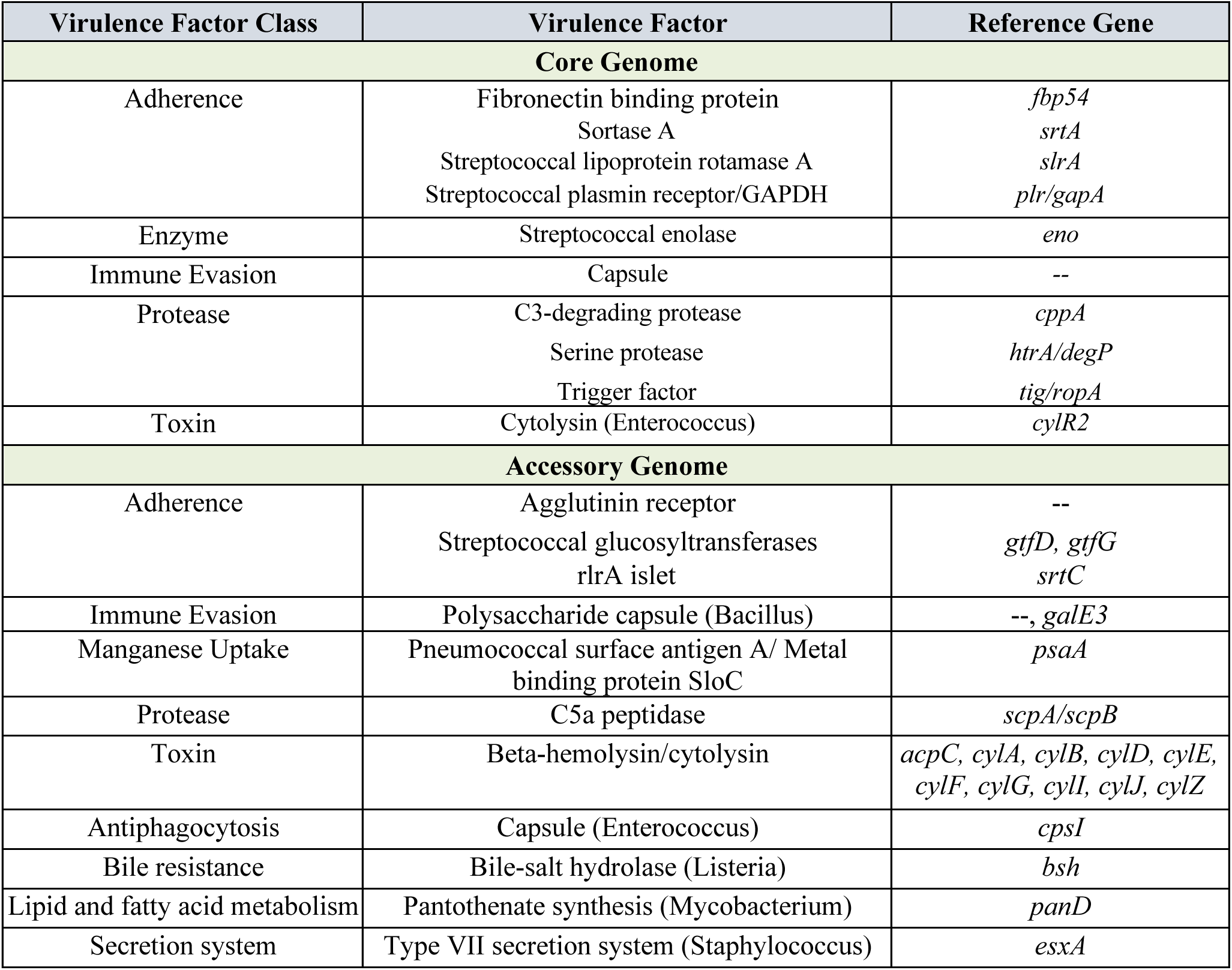
Putative virulence factors in the core and accessory genomes of *Sgg*.

Next, we examined the core and accessory genomes for genes mediating resistance to antimicrobial agents, using the online Resistance Gene Identifier (RGI) (Alcock *et al*., 2023). Two genes involved in vancomycin resistance were identified in the core genome (Supplementary Data, Table S3). The accessory genome contains 18 resistance genes against various classes of antibiotics, including aminoglycosides, lincosamides, macrolides, and tetracyclines (Supplemental Data, Table S3). Consistent with previous findings (Périchon *et al*., 2025), a large proportion of the resistance genes (37%) are involved in resistance to tetracyclines, possibly due to the widespread usage of tetracyclines in the treatment of livestock (Wu *et al*., 2024).

### 3.4 Type VII secretion system locus

To date, functional characterization of T7SS in *Sgg* has been reported in two studies. A T7SS from strain TX20005 was demonstrated to mediate virulence activities *in vitro* and *in vivo* (Taylor *et al*., 2021). TelE, an LXG toxin from strain UCN34, was found to exhibit membrane pore-forming toxicity when expressed in *E. coli* (Teh *et al*., 2023). These studies support the importance of the T7SS in *Sgg* colonization of the host and virulence.

To gain more detailed knowledge of the T7SS, we examined the 76 *Sgg* genomes for the presence of a T7SS locus. To do this, a custom BLAST database containing the 76 genomes was generated and searched using protein sequences of EsxA and EssC, the two well-conserved components of T7SS. The results were then used to locate the T7SS locus on each chromosome and to examine additional genes within the locus. We found that the majority of the strains (67 strains; 88.2%) contain a single T7SS locus encoding all core T7SS components (EsxA, EsaA, EssA, EsaB, EssB, and EssC). A T7SS locus was not detected or was only partially detected in the remaining ten strains. Two of these strains have complete genomes and genuinely lack a T7SS locus. The other eight genomes contain assembly gaps that may account for the apparent absence or incompleteness of the locus. These eight strains were therefore classified as “undetermined” (Supplementary Data, Table S2). Overall, the results indicate that T7SS is widely distributed among *Sgg* strains.

Previous studies demonstrated that the C-terminal region of approximately 200 amino acids of EssC acts as specificity determinant for downstream effector genes (Warne *et al*., 2016; Jäger, Kneuper and Palmer, 2018; Lebeurre *et al*., 2019; Zhou *et al*., 2022; Spencer *et al*., 2023). We therefore examined the EssC C-terminal region in all identified *Sgg* T7SS loci, as well as the genes located downstream of *essC*. We found that all strains containing the previously reported subtype 1 T7SS shared an identical EssC C-terminal region, which was followed by the same downstream gene organization (Fig. 3A). On the other hand, the EssC C-terminal region from the previously described subtype 2 strains showed substantial sequence divergence. We identified four unique C-terminal regions exhibiting amino acid sequence identities from 21.5% to 47.5% to each other (Fig. 3B). Each of these was followed by a distinct set of downstream genes, hence we further classified them into subtype 2A to 2D (Fig. 3A). Interestingly, subtype 2B has a second truncated EssC that shares the same C-terminal region as the EssC in subtype 2A, and is followed by the same set of downstream genes as in 2A (Fig. 3A). This observation suggested that 2A and 2B subtypes might have diverged from a common ancestral subtype through recombination events. We should also point out that the *essC* gene in subtype 2D contains a frameshift mutation, which may result in a non-functional T7SS. In addition to these, we also identified a T7SS locus with a subtype 2A EssC but is followed by a different set of downstream genes and an orphan EssC that is truncated at both the N- and C-terminus. We classified this T7SS locus as subtype 2E (Fig. 3A). Together, our analysis resulted in a total of six T7SS subtypes.

**Fig 3.**
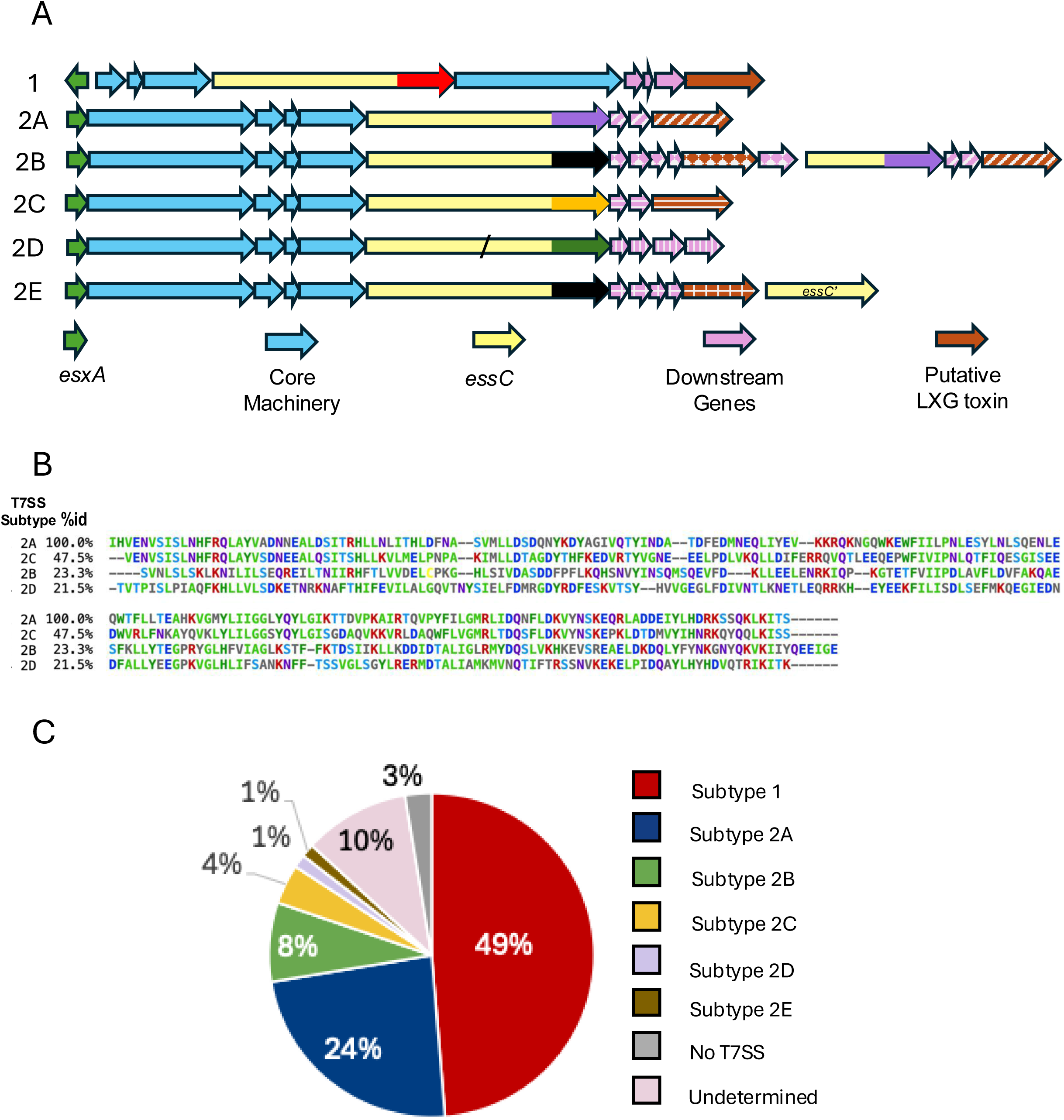
T7SS genetic loci and subtype distribution in *Sgg*. **A.** A schematic diagram of the T7SS locus for subtype 1 to 2E. e*sxA* gene is shown in green arrows, core machinery components *esaA*, *essA*, *esaB* and *essB* are marked in turquoise, and *essC* is shown in yellow with the C-terminal region marked in different colors to indicate the sequence heterogeneity in this region. Downstream genes are shown in pink with patterns except for LXG domain-containing proteins, which are shown in brown with patterns. The different patterns represent sequence heterogeneity in these genes. EssC in subtype 2D has a frameshift mutation which is represented by a slash. **B**. Sequence heterogeneity in the C-terminal region of EssC from subtypes 2A to 2D. The C-terminal 200 amino acid residues of EssCs from representative genomes of the 2A to 2D subtypes were aligned using Clustal Omega and formatted in MView. The amino acid residues are shaded according to identity. The percentage identity to each other over the entire length of the protein (%id) is shown to the left of the alignment. **C.** The prevalence of the six T7SS subtypes among 76 *Sgg* genomes.

We next examined the distribution of the six T7SS subtypes among *Sgg* strains and their relationship to ST, isolation source and geographical region. Subtype 1 is the most prevalent among *Sgg* strains, present in 48.7% (37/76) of all strains (Fig. 3B and Supplementary Data, Table S2).

Among the five newly defined subtypes, 2A is the most prevalent (18/76, 23.7%), followed by 2B (6/76, 7.9%), 2C (3/76, 3.9%), 2D (1/76, 1.3%), and 2E (1/76, 1.3%) (Fig. 3C and Supplementary Data, Table S2). Of the four subtypes that have been found in more than one strain, 2B was detected exclusively in human isolates (Fig. 2 and Supplementary Data, Table S2). Correlation analysis further revealed a significant association between T7SS subtype and isolation source (*p* = 0.0119, Table 2). In addition, we found that T7SS subtype is strongly associated with ST (*p* = 0.0001) (Table 2, Fig. 2, and Supplementary Data, Table S2). Of the four most prevalent STs, ST12 and ST5 strains have subtype 1 T7SS, ST26 has subtype 2A, and ST117 possesses subtype 2B (Fig. 2 and Supplementary Data, Table S2). We also found a significant correlation between T7SS subtype and geographic region (*p* = 0.0279, Table 2). Together, this resulted in new T7SS subtypes identified and correlation statistics suggested that T7SS subtype is strongly associated with *Sgg* lineage.

### 3.5 LXG domain-containing polymorphic toxins

LXG toxins have emerged as one of the most widely distributed classes of T7SSb-dependent effectors and primarily mediate interbacterial competition (Boardman, Palmer and Alcock, 2023).

We identified putative LXG toxin-encoding genes downstream of *essC* in all *Sgg* T7SS subtypes except for 2D (Fig. 3A). Previous studies have described LXG toxin-encoding genes both within and outside the T7SS locus (Whitney *et al*., 2017; Teh *et al*., 2023; Garrett and Palmer, 2024). To obtain a comprehensive inventory of putative LXG toxins, we performed a genome-wide search across the 76 *Sgg* genomes using the protein sequences of characterized LXG toxins from various bacteria as well as the LXG domain consensus sequence from NCBI-CDD, as queries. This search identified a total of 23 unique putative LXG toxins, encoded by genes downstream of *essC* in the T7SS locus and genes offsite. For ease of description, we labeled these 23 toxins as LXG1 – 23 (Fig. 4). These toxins all have the domain arrangement typical of LXG toxins, *i.e.*, an N-terminal LXG domain required for secretion followed by a central and C-terminal domain that presumably mediates toxin activity.

**Fig 4.**
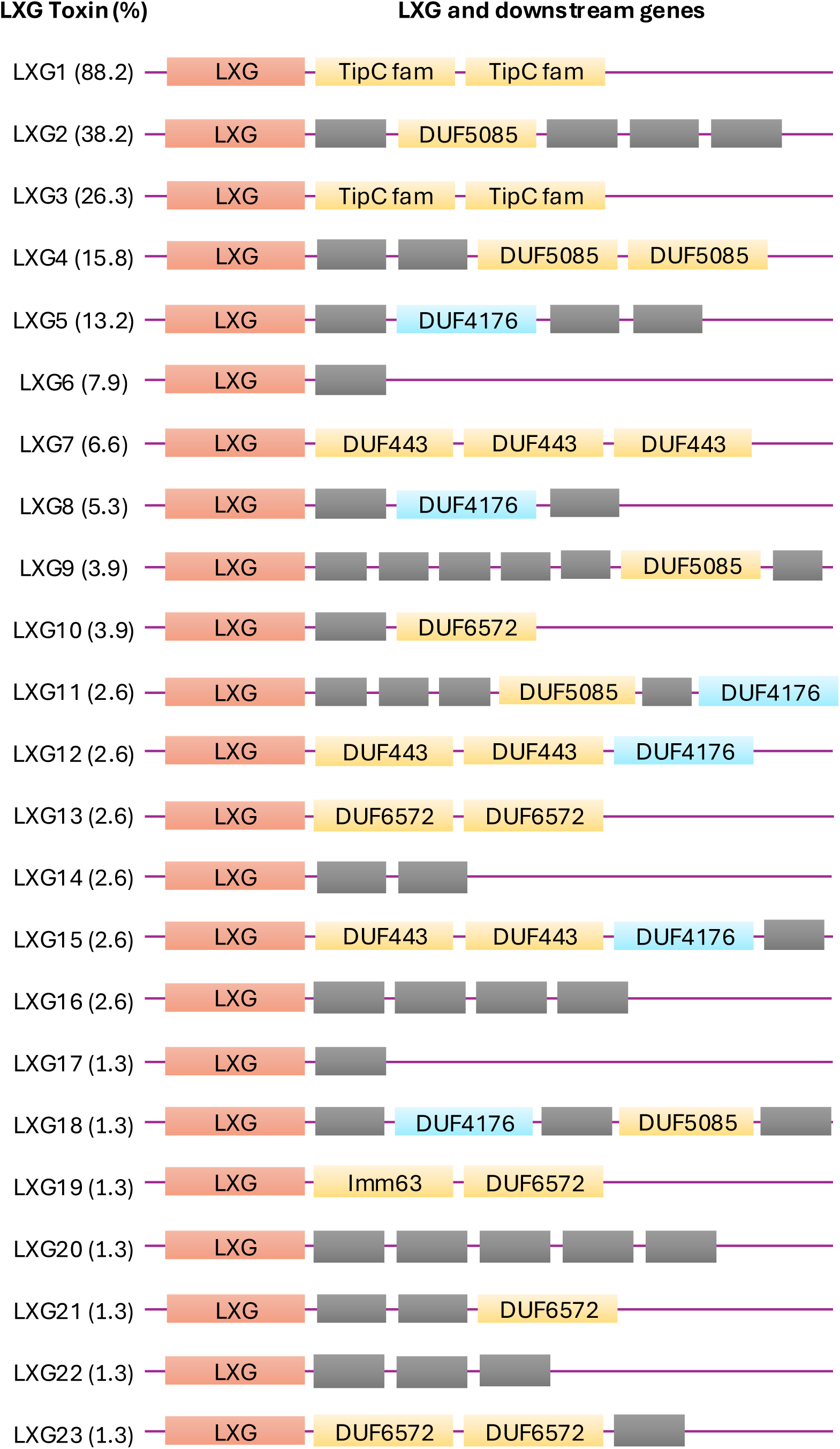
LXG domain-containing polymorphic toxins in *Sgg*. A schematic diagram of the genetic loci of the LXG toxin encoding gene and the downstream genes from representative genomes. LXG toxins were identified by searching the *Sgg* genome database using known LXG toxins and LXG domain consensus sequence as queries, as described in Materials and Methods. Downstream genes were analyzed by searching each gene against the ClusteredNR database at NCBI using Blastp. The prevalence of each LXG toxin as a percentage of total strains is shown to the left of the corresponding genetic cassette in parenthesis. Genes are not drawn to scale with respect to their length.

Among the 23 putative LXG toxins, three were previously reported in *Sgg* strain UCN34 (Teh *et al*., 2023). Specifically, LXG2 corresponds to TelE, whereas LXG1 and LXG3 correspond to GALLO_1068 and GALLO_1574 (Teh *et al*., 2023), respectively (Fig. 4). TelE was characterized as a glycine zipper containing pore-forming toxin. LXG1 and LXG3 share sequence homology (65% and 43% identity, respectively) to TelC from *S. intermedius*, which is a lipid II phosphatase (Whitney *et al*., 2017). Hence LXG1 and LXG3 likely possess a similar enzymatic activity. The rest of the LXG toxins have not been described previously in *Sgg*. Among these, LXG12 exhibits sequence homology (32.7% identity) to TspA, a T7SS-dependent membrane-depolarizing toxin of *Staphylococcus aureus* (Ulhuq *et al*., 2020). To gain further insights into possible functions of the remaining LXG toxins, we searched for protein motifs in their central and C-terminal regions. One protein (LXG19) was found to contain a Tuberculosis necrotizing toxin domain (TNT; PF14021), which acts as an NAD+ glycohydrolase. The previously described TelB of *S. intermedius* (Whitney *et al*., 2017) exhibits NADase activity. Hence, LXG19 may be functionally similar to TelB. We did not identify any motifs in the central or C-terminal regions in the rest of the LXG proteins (E value cut off 0.001) (Fig. 4), suggesting that they are likely novel LXG toxins with uncharacterized functions.

The distribution of the 23 putative LXG toxins in *Sgg* was examined. The top 5 most prevalent LXGs are LXG1 (88.2% of all strains), LXG2 (38.2%), LXG3 (26.3%), LXG4 (15.8%), and LXG5 (13.2%) (Fig. 4, left column). With respect to connection with T7SS subtypes, some LXG toxins exhibit subtype specificity or bias, while others show no apparent subtype preference. For example, LXG2 is identified exclusively in subtype 1 strains, whereas LXG7 and LXG6 are restricted to subtype 2A and 2B, respectively. LXG3 and LXG5 showed preferential association with subtypes 1 and 2B, and subtype 2A, respectively. In contrast, LXG1, the most prevalent LXG toxin, was detected across all six subtypes and is present in all, except two, strains harboring a T7SS locus.

Secretion of LXG toxins requires the assistance of a pair of LXG-associated proteins (Laps), which are small WXG100-like proteins often encoded by genes located upstream of cognate LXG toxin genes (Klein *et al*., 2022). We investigated the upstream genes of LXG1 in selected strains from each T7SS subtype. We found two genes encoding small proteins of 94 and 114 amino acids, respectively, upstream of the LXG1 gene in each of the select genomes. The corresponding sequences between the different subtypes are nearly identical (> 96% identity over the entire length of the proteins). Analysis of the two protein sequences showed that they are DUF3130 and DUF3958 domain-containing proteins, respectively, corresponding to LapC1 and LapC2 for TelC in *S. intermedius* (Klein *et al*., 2022). Thus, LXG1, along with its associated Lap proteins, appears to be highly conserved among *Sgg* strains. The presence of this Lap-toxin module in strains carrying different T7SS subtypes raises the possibility that LXG1 may be secreted by multiple T7SS variants. Further investigation will be required to determine whether this is the case.

LXG toxin genes are commonly followed by genes encoding immunity proteins to protect from self-damage. The genes downstream of the putative LXG toxin genes were examined. We found genes encoding proteins with significant sequence homology to previously reported immunity proteins against LXG toxins, including TipC (against TelC) (Klein *et al*., 2018), DUF5085 domain-containing protein (against TelE and EsxX) (Alcock *et al*., 2025), and DUF443 domain-containing proteins (against TspA) (Ulhuq *et al*., 2020; Yang *et al*., 2024), downstream of multiple LXG toxin genes (Fig. 4). In addition, we also identified proteins with similarity to Imm63 family proteins and DUF6572 domain-containing proteins encoded by genes downstream of several LXG toxin genes (Fig. 4). These protein families have been described as immunity proteins for toxins in Gram-negative bacteria and for T7SS-dependent MSE-ExT toxins, respectively (Zhang *et al*., 2012; Yang *et al*., 2023; Bai *et al*., 2025). Recent studies indicated that DUF4176 domain-containing proteins act as molecular chaperones that stabilize the LXG toxins for secretion (Gkragkopoulou *et al*., 2025; Wu *et al*., 2025). We found DUF4176 domain-containing proteins downstream of multiple LXG toxins (Fig. 4). Altogether, 17 out of the 23 putative toxin genes are followed by genes encoding potential immunity proteins or a DUF4176 domain-containing chaperone in close proximity.

### 3.6 The ability to form biofilms correlates with ST and T7SS subtype

The ability to form biofilms is an important virulence trait and a contributing factor to treatment failure. We evaluated the biofilm formation capacity among the strains in the lab collection using a standard crystal violet assay. Based on the mean values from three independent experiments, strains were categorized into three groups according to their relative biofilm formation capacity (Fig. 5A): strong biofilm formers (≥75th percentile, blue bars), intermediate biofilm formers (25th–75th percentile, orange bars), and weak biofilm formers (≤25th percentile, red bars). Correlation analysis revealed that biofilm formation under the experimental conditions used here is significantly correlated with ST (*p* = 0.0262) and T7SS subtype (*p* = 0.0059) (Table 2, Fig. 2 and Fig. 5B). With respect to T7SS subtypes, eight of the nine (88.9%) strong biofilm-forming strains have subtype 1 T7SS, and the last one has no T7SS (Fig. 5B). In contrast, none of the seven weak biofilm-forming strains have subtype 1 T7SS. Instead, three of them have subtype 2A (42.9%), three have subtype 2B (42.9%), and the last one has an undetermined subtype (Fig. 5B). These results highlight a possible previously unrecognized association between T7SS subtype and biofilm formation capacity in *Sgg*.

**Fig 5.**
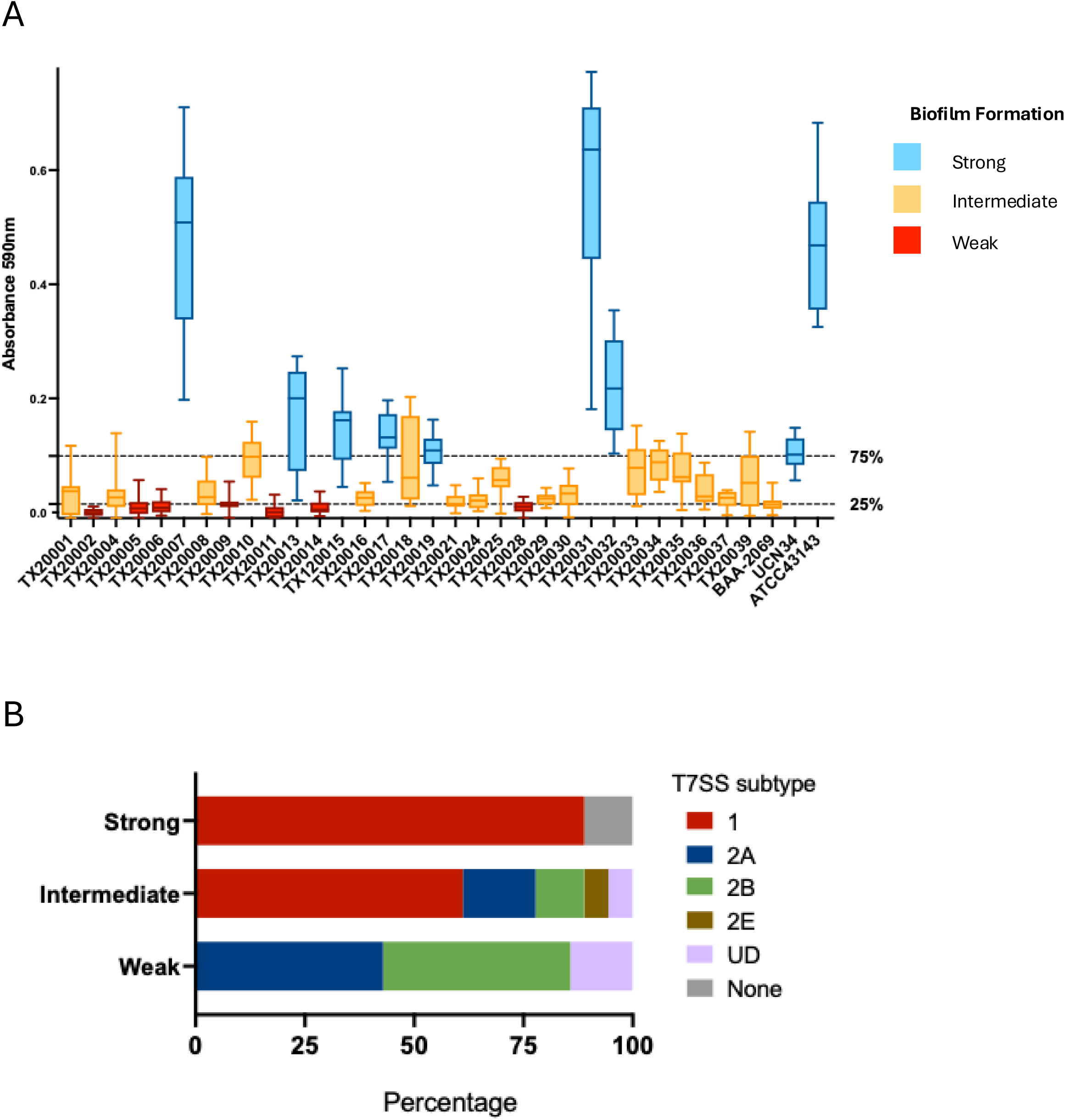
Biofilm formation capability of *Sgg* strains. **A.** Overnight cultures of *Sgg* strains were diluted to OD600nm of 0.1 in fresh BHI broth and incubated in 96-well plates for 24 hours, as described in Materials and Methods. Biofilm formation was measured using crystal violet staining. The mean absorbance from three independent experiments (each with 3 technical replicates) was used to categorize the strains into three groups: strong (blue, above 75^th^ percentile), intermediate (orange, between 75^th^ and 25^th^ percentile), and weak (red, below 25^th^ percentile mark) biofilm formers. The dashed lines indicate the 75^th^ (top) and 25^th^ (bottom) values, respectively. The graph is a box and whiskers plot with the median value shown as a line in the box and whiskers showing the min to max values. **B.** The distribution of T7SS subtypes in the strong, intermediate and weak biofilm-forming groups.

Whether subtype 1 T7SS is functionally involved in biofilm formation is unknown. Further studies will be required to elucidate the nature and molecular basis of these associations.

### 3.7 The ability to promote colon cancer cell proliferation

Previous studies showed that *Sgg* strains displayed varied abilities to promote colon cancer cell proliferation (Kumar *et al*., 2017, 2018; Taddese *et al*., 2020). We further investigated the ability of 34 *Sgg* strains in the lab collection to promote the proliferation of human colon cancer cell lines HT29 and HCT116. The cells were co-cultured with each *Sgg* strain for a total of 48 hours and cell proliferation was determined using the CCK-8 viability assay. To control for absorbance from bacteria, *Sgg* suspensions were added to wells with no cells in parallel and were subject to the same procedure. Absorbance from the bacteria-only samples was then subtracted from the corresponding co-culture samples. Of the 34 strains, nine (26.5%) statistically increased cell proliferation in both cell lines (Fig. 6A – 6D, blue bars), whereas 19 (55.9%) did not (Fig. 6A-6D, red bars). For a few strains, the absorbance from bacteria-only control was higher than that from their corresponding co-culture samples, resulting in negative values after subtraction (Fig. 6A – 6D, orange bars). TX20005 was previously shown to stimulate the proliferation of HT129 and HCT116 cells, making our results consistent with these findings (Kumar *et al*., 2017). One strain (TX20018) showed a pro-proliferative phenotype in HCT116 (Fig. 6A) but not in HT29 (Fig. 6C), suggesting a cell-line-specific effect of this strain. Correlation analysis showed no statistically significant association between the ability of *Sgg* strains to stimulate cell proliferation with ST or T7SS subtype (Table 2). These results suggest that the pro-proliferative phenotype may be mediated by multiple and strain-specific mechanisms. A previous study comparing *Sgg* strains from CRC patients versus non-cancer individuals did not identify virulence pathways specific to CRC-associated isolates (Périchon *et al*., 2025). Our results are consistent with this earlier observation.

**Fig 6.**
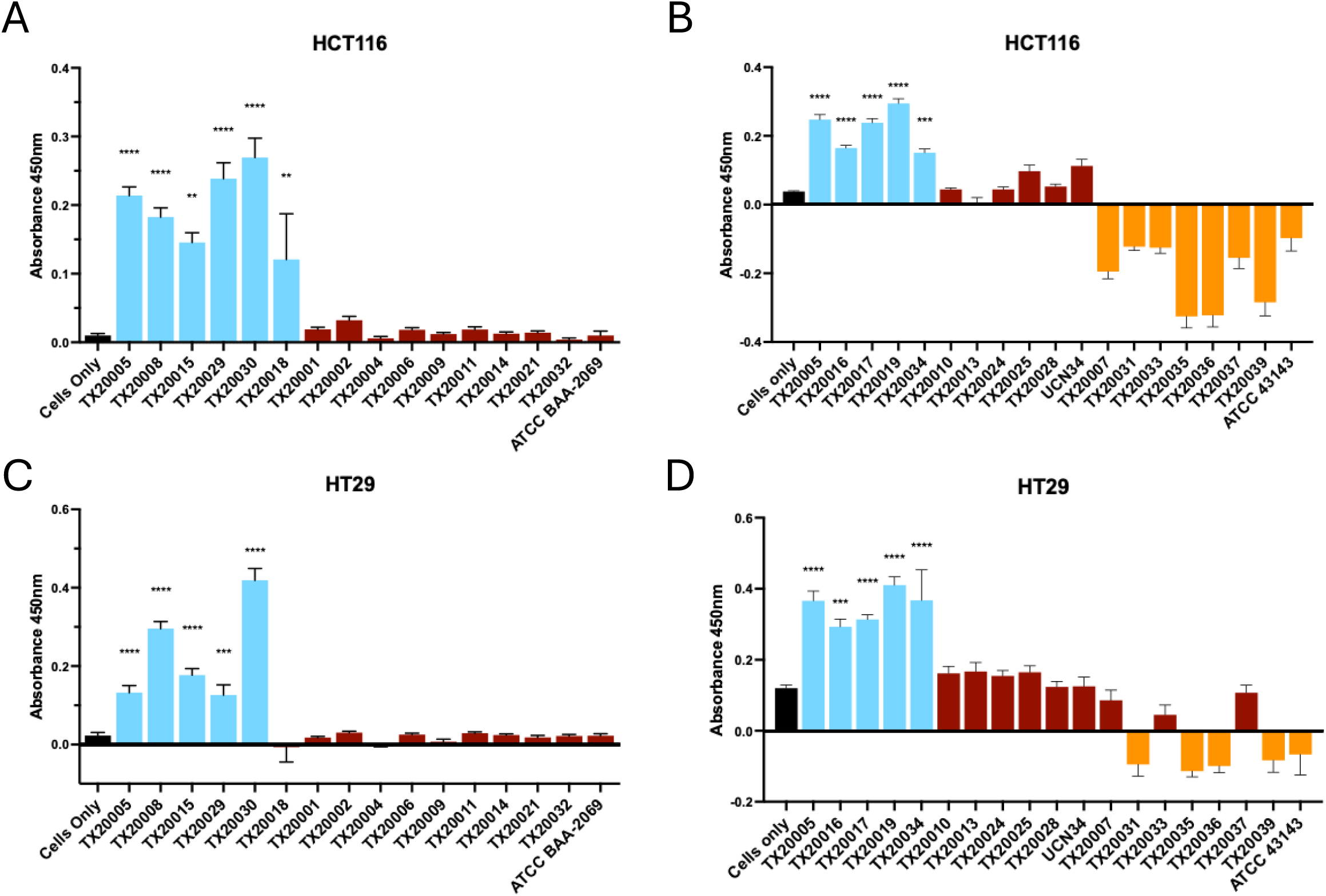
The ability of *Sgg* to stimulate CRC cell proliferation is strain specific. Human CRC cell lines HCT116 (**A** and **B**) and HT29 (**C** and **D**) were co-cultured with *Sgg* strains at an MOI of 1 in 96-well plates. For each *Sgg* strain, bacterial suspensions were also added to wells without cells in parallel to obtain readings for the bacterial background. Cell proliferation was measured using CCK-8, as described in Materials and Methods. The absorbance values shown in the graph were after subtraction of absorbance from media only and bacterial only from the corresponding strain. Bars are colored based on their phenotype: blue, pro-proliferation; red, non-pro-proliferation; and orange, bacteria only absorbance higher than co-culture absorbance. Data shown is mean ± standard error of the mean, combined from at least three independent experiments, each with 3 technical replicates. Statistical analysis was performed using a one-way ANOVA with Dunnett’s multiple comparison test to compare results for each strain vs. cells only. ****, *p* < 0.0001; ***, *p* < 0.001; **, *p* < 0.01.

## 4 Discussion and conclusions

In this study, we performed a comprehensive genomic analysis of 76 *Sgg* strains isolated from human and animal sources at diverse geographical locations across four continents. Our analysis identified dominant STs that were significantly correlated with geographical location. Several STs are restricted to human isolates in our dataset. It should be noted that the number of environmental isolates in our data set is limited and studies involving larger cohorts will be needed to determine whether the observed association pattern extends beyond the strains analyzed here. Analysis of putative virulence factors in both the core and accessory genomes also suggests that host adherence and immune evasion are two central features of the *Sgg* pathobiont lifestyle.

Our systematic analysis of the T7SS loci in the 76 genomes revealed that T7SS is widely distributed among *Sgg* strains. In addition, our analyses uncovered five new T7SS subtypes, underscoring the genetic complexity of T7SS in *Sgg*. Furthermore, the identification of 23 unique putative LXG toxins substantially expands the current repertoire of LXG toxins in *Sgg*. The LXG toxins have emerged as one of the most widely distributed effector classes secreted by T7SSb in Firmicutes; however, knowledge of their biological activities remains limited. Notably, many of the 23 putative LXG toxins identified here do not contain recognizable functional motifs, further highlighting the heterogenous nature of this toxin family and the need for additional studies to elucidate their biological roles. Interestingly, we observed distinct distribution patterns among LXG toxins, including subtype-restricted, subtype-biased, and broadly distributed (pan-subtype) toxins. Future studies will be required to understand the interactions between these different types of LXG toxins and the secretion machinery of specific T7SS subtypes.

Our work also identified a significant correlation between *Sgg* T7SS subtype and the capacity of the bacteria to form biofilms. Specifically, strains possessing subtype 1 T7SS tend to display strong or intermediate biofilm-forming capacity and are absent from the weak biofilm-forming group. Grimm and colleagues reported that *Sgg* exhibits strain-dependent biofilm formation in BHI media and a strain-dependent response to the effect of lysozyme on biofilm formation (Grimm *et al*., 2018). Teh and colleagues showed that biofilm formation in *Sgg* strain UCN34 is regulated by intracellular cyclic di-AMP levels (Teh *et al*., 2019). To our knowledge, a connection between T7SS and biofilm formation has not been previously reported in *Sgg*. Whether this connection reflects a causal relationship between *Sgg* subtype 1 and enhanced biofilm formation capacity remains unclear. In other bacteria, including mycobacteria, *Staphylococcus aureus*, and *Bacillus subtilis,* T7SS has been implicated in biofilm formation (Lai *et al*., 2018; Nath, Ray and Buragohain, 2018; Pena *et al*., 2019; Finn, Luzinski and Burton, 2024). Although the molecular basis underlying this involvement are not fully understood, proposed mechanisms include cell wall alterations, sporulation, and quorum sensing. Further investigations will be required to determine whether T7SS plays a functional role in *Sgg* biofilm formation.

In summary, our work advances understanding of the genomic landscape and T7SS heterogeneity in *Sgg*, revealing lineage-associated patterns and a previously unrecognized link to biofilm formation. While the molecular basis of these associations remains to be further characterized, our findings underscore the complexity of T7SS-mediated functions in this pathobiont. Future studies will be required to clearly define how the different T7SS subtypes contribute to colonization, competition, and virulence.

## 5 Conflict of Interest

The authors declare that the research was conducted in the absence of any commercial or financial relationships that could be construed as a potential conflict of interest.

## 6 Author Contributions

GC, YX, SW and JT were involved in the acquisition of experimental data. GC, IP, and YX contributed to methodology and data analysis. YX and GC drafted the manuscript. YX and JH provided resources and supervision.

## 7 Funding

This study was supported by the National Institute of Allergy and Infectious Diseases awards R01AI63206 and R21AI166688 to Y. Xu and 3R01AI63206-01A1S1 to G. Calderon.

## Acknowledgments

We thank Amarachukwu Okeke, University of Houston, Houston, TX, for assistance in the cell proliferation assays and Dr. Jessica Galloway-Peña, Texas A&M University, College Station, TX, for providing constructive suggestions for genomic analysis. ChatGPT (OpenAI) was used to improve the clarity and grammar of portions of the manuscript. The authors reviewed and edited all the outputs.

## Notes

### Competing Interest Statement

The authors have declared no competing interest.

